# A 3D-printed and freely available device to measure the zebrafish optokinetic response before and after injury

**DOI:** 10.1101/2023.08.15.553448

**Authors:** Ashley Hermans, Sophia Tajnai, Allison Tieman, Sarah Young, Ashley Franklin, Mackenzie Horutz, Steven Henle

## Abstract

Zebrafish have eyes similar to humans, making them a beneficial model organism for studying vision. However, zebrafish are different in that they can regenerate their optic nerve after injury, which most other animals cannot. Measuring vision in people who have communicative ability is achieved using eye charts. Because fish cannot use an eye chart, we utilize the optokinetic response (OKR) that is present in virtually all vertebrates to determine if a zebrafish has eyesight. To this end, we have developed an inexpensive OKR setup that uses 3D-printed and off-the-shelf parts. This setup has been designed and used by undergraduate researchers and is also scalable to a classroom lab setup. We demonstrate that this setup is fully functional for assessing OKR, and we use it to illustrate the return of the OKR following optic nerve injury in adult zebrafish.

## Introduction

Zebrafish are an excellent model for studying vision because they have all the advantages of other small vertebrate models, while retaining many anatomical similarities to humans^1,2^. A key difference in the zebrafish visual system is that adult zebrafish can regenerate their optic nerve after injury,^3,4^ whereas humans lack this ability. However, unlike humans, zebrafish can’t perform a standard vision test, which makes assessing visual functions a bit more challenging. Researchers use several different visual tests to evaluate visual function in zebrafish. The optokinetic response assay (OKR) is one of the most commonly used.^5,6^ The optokinetic response is an innate visual response performed by most vertebrates in response to a moving object (think of a train passing by); their eyes will track the movement and then snap back in a saccade and continue tracking the movement.

Traditionally, to measure the OKR, the animal is restrained and placed in the middle of a rotating drum or surrounded by screens, which display moving bars. Video is then captured from above, often through a stereomicroscope. Though the setup is not necessarily that complicated, commercially available devices can cost more than $40,000, and previously published homemade versions require the use of a machine shop.^7^ Neither of these options is ideal for educational settings. Here we present a 3D printable and open-source OKR setup that can be made for around $50. We demonstrate this setup can measure the optokinetic response in adult zebrafish across a range of speeds.

Additionally, we are interested in analyzing the recovery of visual function following optic nerve injury in adult zebrafish. Optic nerve regeneration is traditionally measured by analyzing the length of optic nerve regrowth.^8^ This requires euthanizing the animal to perform pathological analysis, only allows you to measure one timepoint per animal, and does not provide information about the recovery of visual function. Here we show that our OKR device can measure functional visual recovery of the optokinetic response in an undergraduate setting. We hope that the ease, cost, and scalability of this OKR setup will promote its use in educational research and classroom settings.

## Methods

### Construction of the OKR Device

The OKR device consists of four 3D-printed parts, a stepper motor and controller, a belt and pulley, and an Arduino controller (Table 1). Arduino controllers are easy-to-program microcontrollers that give you more systematic control over the OKR speed and make it more adaptable. The code to run the motor for the OKR is provided (Supplemental Data 5). However, if you are interested in customizing the setup or learning more, an excellent place to start is the Arduino documentation page on stepper motors (https://docs.arduino.cc/learn/electronics/stepper-motors).

To assemble the device, first, print the four 3D printed pieces. PLA plastic is sufficient and easy to print with. The drum rests in the drum base, and then the bottom motor holds rest against the drum base (Fig. 1A). Place the motor into the top motor holder with the pulley. Wrap the belt around the drum and pulley and place the motor setup into the bottom motor holder.

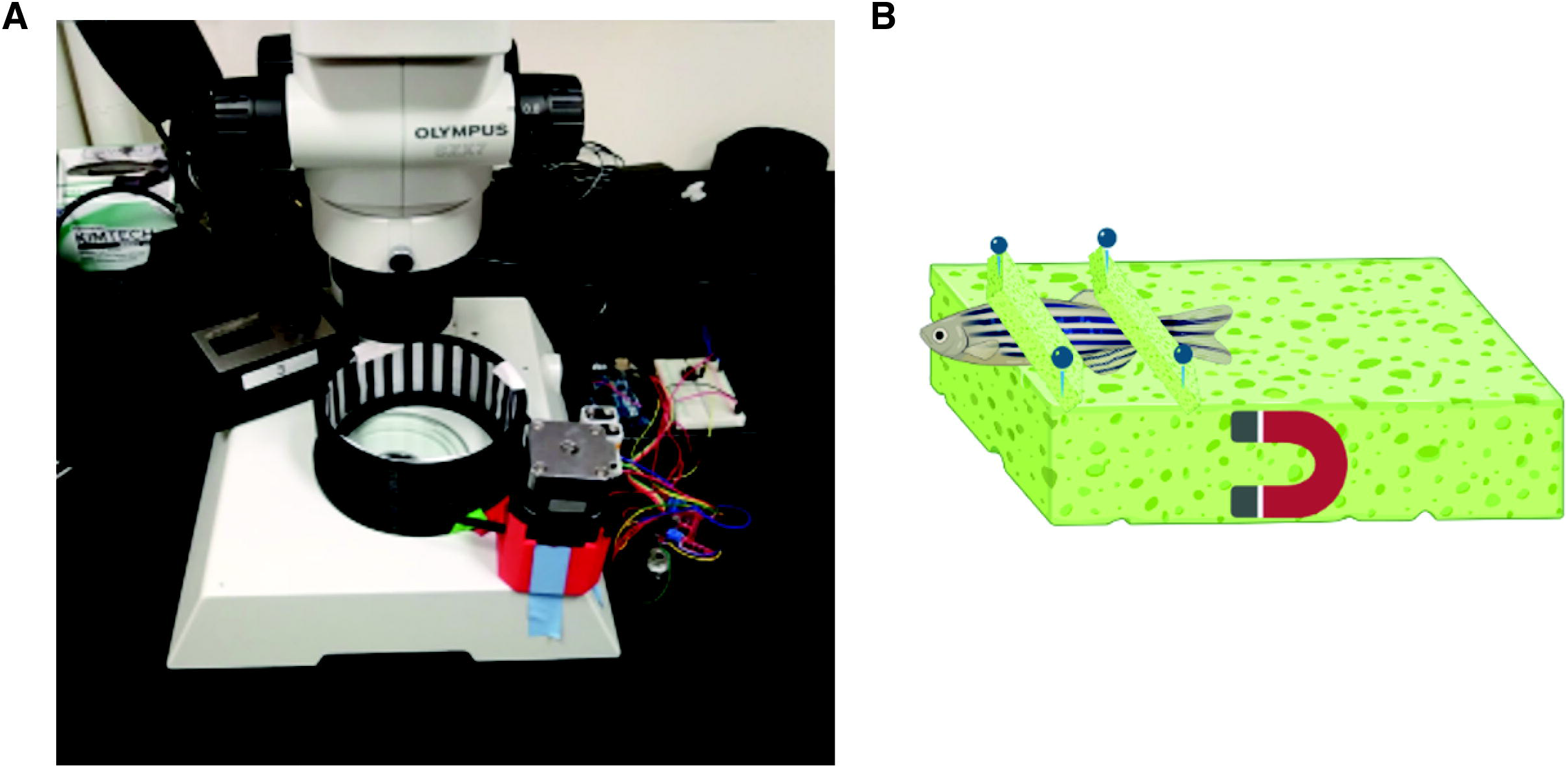

Connect the motor to the motor controller and Arduino controller as shown here: https://docs.arduino.cc/learn/electronics/stepper-motors. One to two 9V batteries are needed depending on the motor used. You can also connect a computer to a USB port to help power the motor if preferred.

The drum has a slot all the way around it to fit a piece of paper with patterns of bars. You can experiment with bar size, contrast, and color easily this way by printing out different size sheets of paper. For standard assays, we use max contrast ½” wide bars. We recommend that you calibrate your OKR setup by taking a video of you on a rotating piece of paper with one mark on it for each selected speed. You can then review the video to calculate revolutions per minute (rpm).

If OKR analysis is performed on a fish with an injured optic nerve, the uninjured eye is blocked from seeing the stimulus by placing a solid-colored piece of paper in front of the appropriate visual field for that eye. The stimulus is then just shown to the injured eye, but the movement is only measured in the uninjured eye. This is because the optic nerve injury procedure often injures ocular muscles affecting eye movement, but since the OKR will happen in both eyes, even if just one eye can see it, we can test vision from the injured eye by monitoring movement in the uninjured.

### Performing the OKR Assay

To perform an OKR test, the fish is first anesthetized using 500ml/l tricaine for no less than 30 seconds and no more than 1 minute. It is removed from the tricaine solution with a slotted spoon five seconds after gill movement has ceased. The fish is placed onto a moist sponge; two small sponges are pinned to the larger sponge to immobilize the fish (Fig. 1B). The bottom of the big sponge is magnetic and will stick to the cylindrical water-filled tank when placed inside. Once the fish is secured in the tank and situated under the microscope, start recording from the camera, and start the motor program through the Arduino controller. This will cause the black and white stripes to rotate around. We used a Flir Blackly camera attached to the camera port of a stereoscope to capture images. However, most camera phones held up to the eyepiece by hand or using a phone camera adaptor (such as Carson Universal Optics Digiscoping Adaptor - https://a.co/d/dspEY1f) can work too, and some phone camera macro lens adaptors may be sufficient as well.

### Analysis of OKR

Videos of the OKR were analyzed using ImageJ (NIH) software, which is freely available here: https://imagej.net/software/fiji/. The video is loaded into the software, and then a line is drawn down the midline of the fish to serve as a reference. Using the angle tool in ImageJ, a point is made on both sides of the eye where the lens is seen to intersect the rest of the eye tissue, and then a third point is clicked, making a perpendicular angle to the midline reference (Fig. 2).

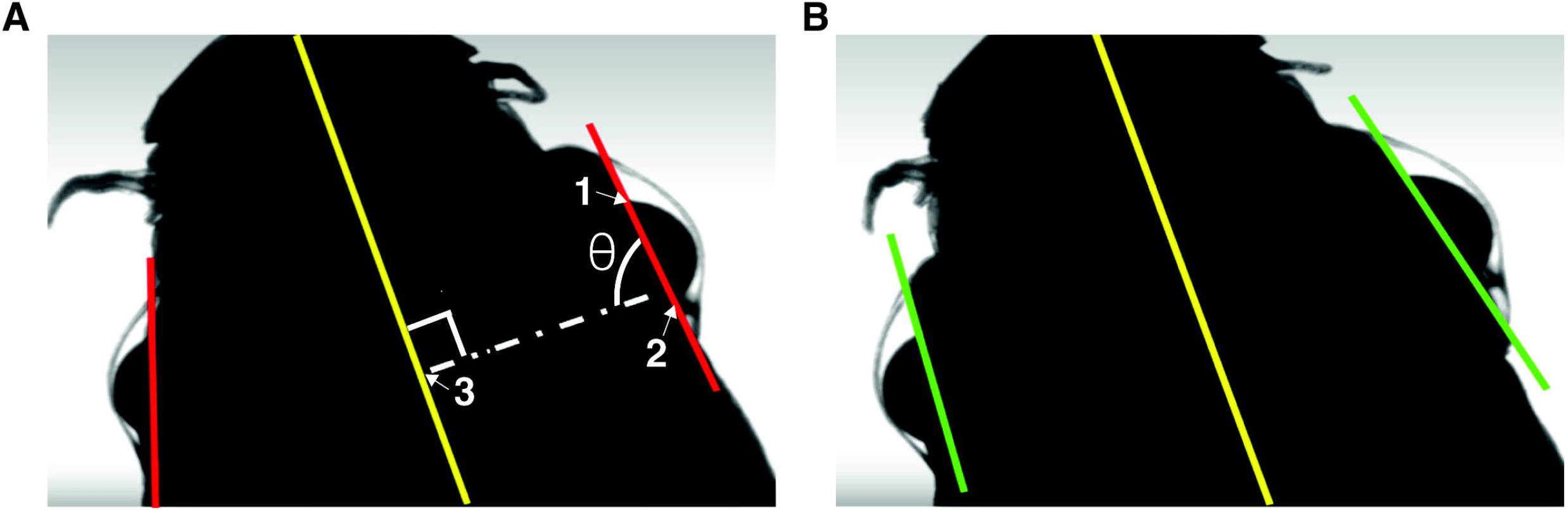

For a quicker overall but less quantitative result, we also grade the videos on a 0-3 scale, with 0 showing no movement, 1 showing eye movements that don’t coordinate with the OKR response, 2 showing limited movements that correlate with the expected OKR response, and 3 demonstrating normal OKR movements.

### Optic Nerve Crush

The optic nerve crush operation is performed by a trained and Citi-training certified researcher, following all IACUC guidelines similar to previously published work^8,9^. The adult zebrafish is anesthetized in a tricaine 500ml/L solution until movement has ceased and the fish does not respond to tapping stimuli on its tail. It is then transferred to a moist sponge. Forceps are used to gently pull the eye away from the socket enough to access the optic nerve. The optic nerve is crushed by releasing reverse forceps which clamp on the optic nerve for 3 seconds. The optic nerve crush is only performed on one eye to spare visual function allowing the fish to continue to thrive during recovery. The Zebrafish is then transferred into a recovery tank using a slotted spoon and monitored until it swims on its own. The fish is monitored throughout the procedure to ensure minimal distress. Fish are allowed at least five days of recovery before performing the OKR assay.

### Zebrafish Care

All procedures using zebrafish were approved by the Carthage College IACUC. All researchers and students handling adult fish completed an occupational health questionnaire and CITI training related to the research.

## Results

We used the described OKR setup to first perform OKR analysis on wildtype uninjured zebrafish. Using the Arduino controller, it is straightforward to systematically increase the revolutions per minute throughout the analysis in a standardized way. We used this to show that adult zebrafish in our OKR setup demonstrates the similar “saw-tooth” pattern of eye movements (Fig. 3, Supplemental Movie S1) that others have reported in their OKR analysis across a variety of speeds.^5,6^ Additionally, this shows that at higher speeds, the OKR actually switches to a slower response pattern (Fig. 3).

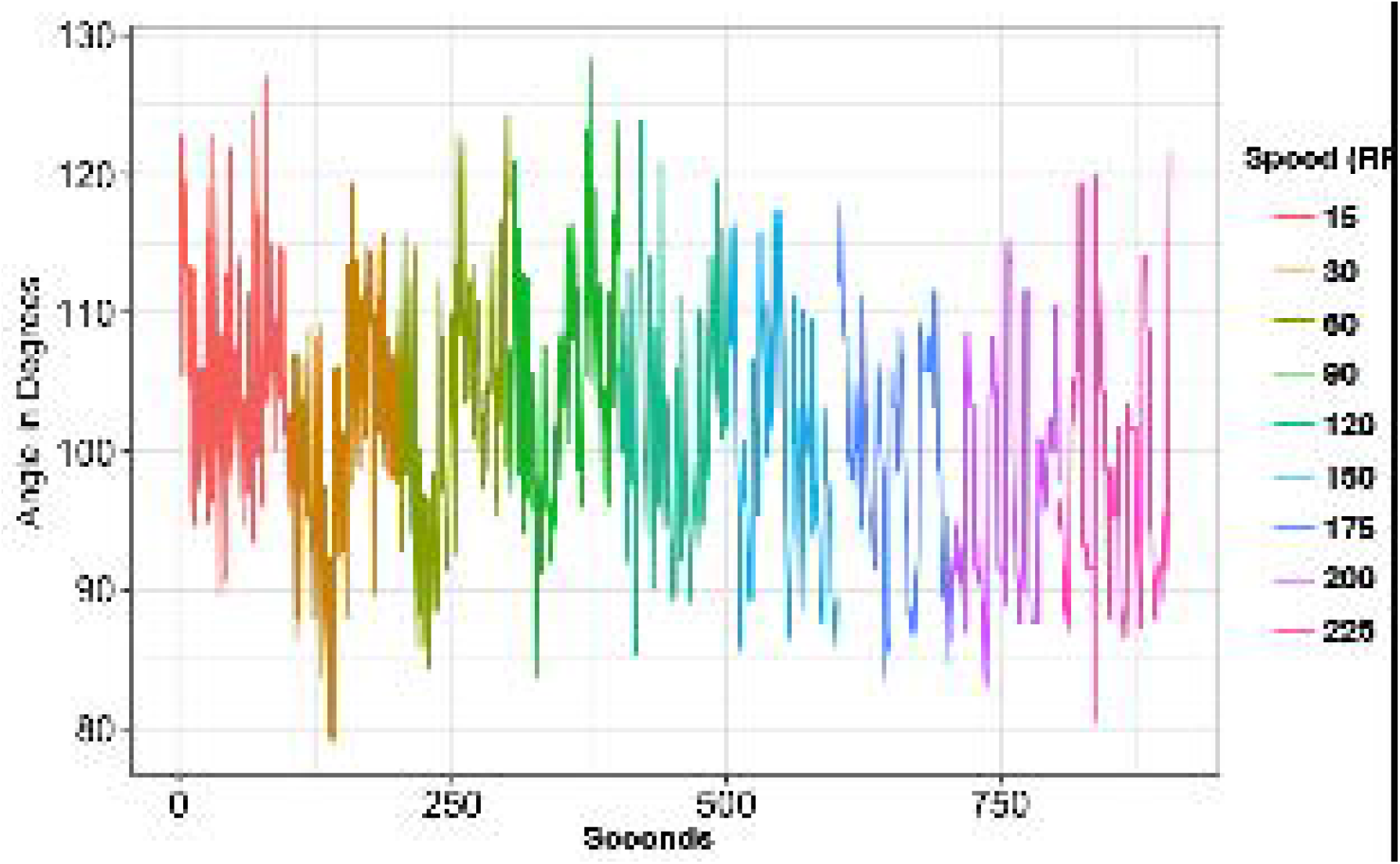

We next performed OKR analysis on adult zebrafish recovering from an optic nerve injury (Fig. 4). These fish showed no visual response at day 5 (the first day measured) and then usually demonstrated full recovery after two weeks. This is in line with previously published anatomical data looking at regenerative optic nerve growth in adult fish.^8^ This demonstrates recovery of visual function occurs nearly as quickly as the rate of axonal growth.

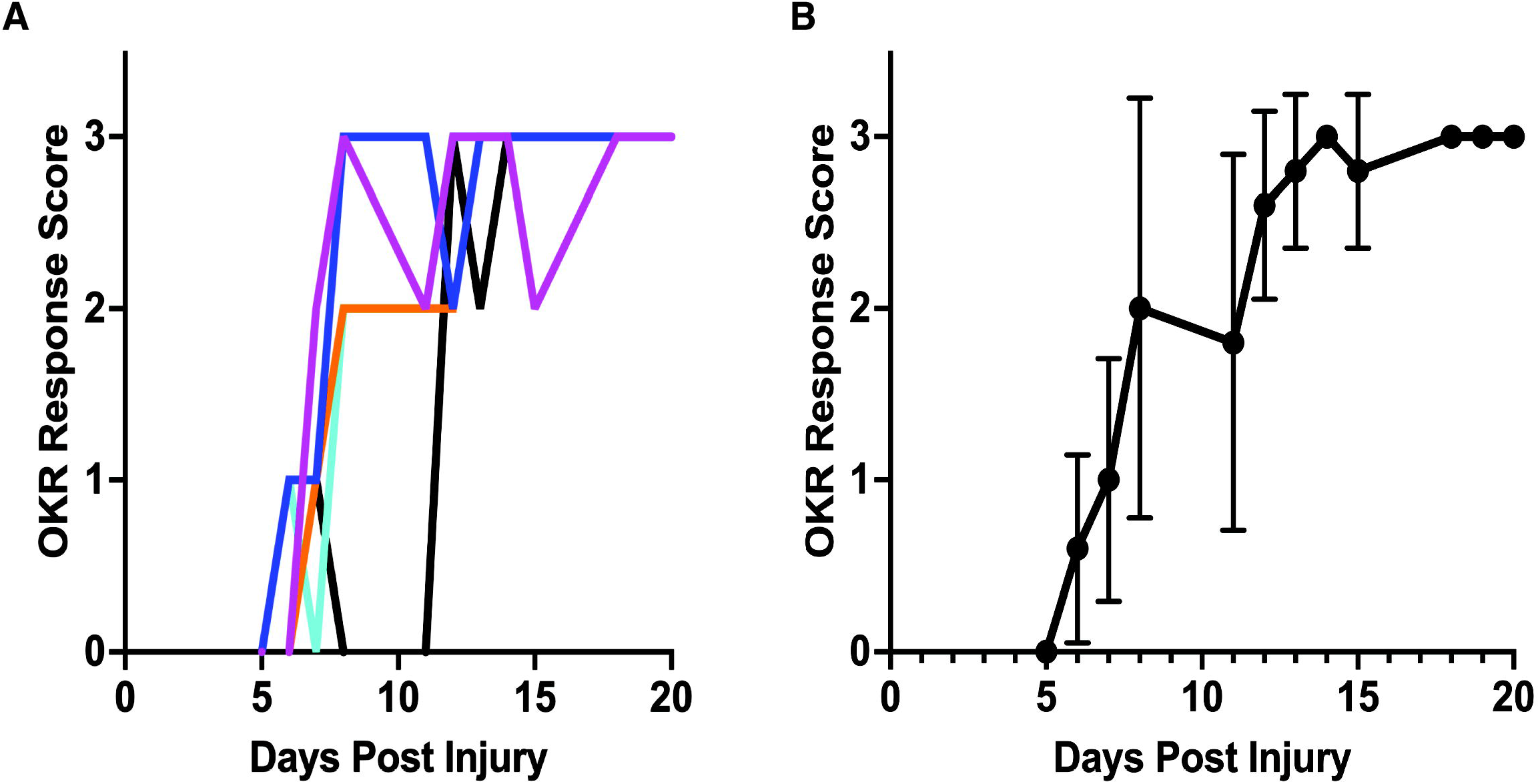

## Discussion

Zebrafish are an excellent model system for studying vision and are great for undergraduate research and teaching. To bring this together, we developed an OKR system that uses a combination of 3D printed and off-the-shelf parts to allow students to build and use a device to study vision in zebrafish. Our device is relatively inexpensive, making it scalable across a classroom lab where groups of students could each have their own setup.

We demonstrated that our OKR setup is effective in eliciting and measuring the OKR in adult zebrafish. Though we haven’t tried it ourselves, we expect it should work for larval zebrafish as well since OKR analysis is more commonly performed on larval fish^5,6^. It is also easy to adapt parameters such as speed and duration through the Arduino controller, or by changing the displayed visual stimuli by printout different patterned bars. We hope that this serves as a sandbox to allow students to devise their own experiments and test out different variables.

In the future, we plan to work to automate the analysis end of the experiment. Currently, manually annotating is the most laborious part of the experiment. In larval zebrafish, software programs exist to automate the analysis^10,11^. However, these tools do not work well with adult zebrafish primarily because the pigmentation difference between the adult eye and the skin is not as great as in the larvae. We are looking to see if AI-based analysis software, such as DeepLabCut^12^, can be trained to overcome this issue.

We further hope that in the future, the scalability of this setup will enable us and others to screen through chemical and genetic mechanisms involved in blinding and regeneration.

## Supporting information

all supplemental files

