## Supplementary material for "A 3D-printed and freely available device to measure the zebrafish optokinetic response before and after injury": all supplemental files: Suppplemental data 5 - Arduino code for OKR motor.ino

```
#include <Stepper.h>

const int button1Pin = 2;
const int stepIN1Pin = 8;         
const int stepIN2Pin = 9;
const int stepIN3Pin = 10;
const int stepIN4Pin = 11;
const int stepsPerRevolution = 200;  // change this to fit the number of steps per revolution
// for your motor

// initialize the stepper library on pins 8 through 11:
Stepper myStepper(stepsPerRevolution, 8, 9, 10, 11);

int stepCount = 0;

void setup() {
 // pins of where the button and motor connect on the uno
 pinMode(button1Pin, INPUT);
 pinMode(stepIN1Pin, OUTPUT);
 pinMode(stepIN2Pin, OUTPUT);
 pinMode(stepIN3Pin, OUTPUT);
 pinMode(stepIN4Pin, OUTPUT);
 
}

void loop () {
 int button1State;
 button1State = digitalRead(button1Pin);
 if(button1State == LOW){   // LOW is when the button is pressed 
        myStepper.setSpeed(15); // in revolutions per a minute
        myStepper.step(3000); // setSpeed*200 to find the 1 minute number
    myStepper.setSpeed(30);
    myStepper.step(6000);
          myStepper.setSpeed(60);
          myStepper.step(12000);
                myStepper.setSpeed(90);
                myStepper.step(18000);
                  myStepper.setSpeed(120);
                  myStepper.step(24000);
                      myStepper.setSpeed(150);
                      myStepper.step(30000);
                     
} //should take 6 minutes from button press to go through the cycle and stop
}
```
